# A 96well ultrafiltration approach for the high-throughput proteome analysis of extracellular vesicles isolated from conditioned medium

**DOI:** 10.1101/2025.01.24.634668

**Authors:** Jarne Pauwels, Tessa Van de Steene, Freya De Muyer, Danaë De Pauw, Femke Baeke, Sven Eyckerman, Kris Gevaert

**Affiliations:** VIB-UGent Center for Medical Biotechnology, VIB, 9052 Ghent, Belgium; Department of Biomolecular Medicine, Ghent University, 9052 Ghent, Belgium; Ghent University Expertise Center for Transmission Electron Microscopy and VIB BioImaging Core, 9000 Ghent, Belgium; Department of Biomedical Molecular Biology, Ghent University, VIB Center for Inflammation Research, 9052 Ghent, Belgium

**Keywords:** 300 kDa MWCO 96well ultrafiltration, TWEEN-20, proteomics, conditioned medium

## Abstract

1.

Extracellular vesicles (EVs), nanoscale vesicles that are secreted by cells, are critical mediators of intercellular communication and play a crucial role in diverse pathologies such as cancer development. Therefore, EVs are regarded as having high potential in the clinic, both for diagnostic and therapeutic applications. Unfortunately, EVs reside in complex biofluids and their consistent isolation at sufficient purity for mass spectrometry-based proteomics has proven to be challenging, especially when increased high-throughput is required. Here, we describe the incorporation of our previously reported filter-aided EV enrichment (FAEVEr) strategy for the isolation of EVs from conditioned medium, from harvest to proteomic analysis completely to a streamlined 96well format. We compared our approach with ultracentrifugation, the most widely used method for EV enrichment, in terms of protein identifications, consistency, reproducibility and overall performance, including the invested time, resources and required expertise. In addition, our results show that including relative high percentages of TWEEN-20, a mild detergent, markedly improves the final purity of the EV proteome by removing the bulk of non-EV proteins (e.g. serum proteins) and significantly increases the number of identified transmembrane proteins. Moreover, our FAEVEr 96well strategy improves the overall reproducibility with a consistent number of protein identifications and decreased number of missing values across replicates. This promotes the validity and comparability between results, which is essential in both a clinical and research setting, where consistency is paramount.

**Graphical abstract:** 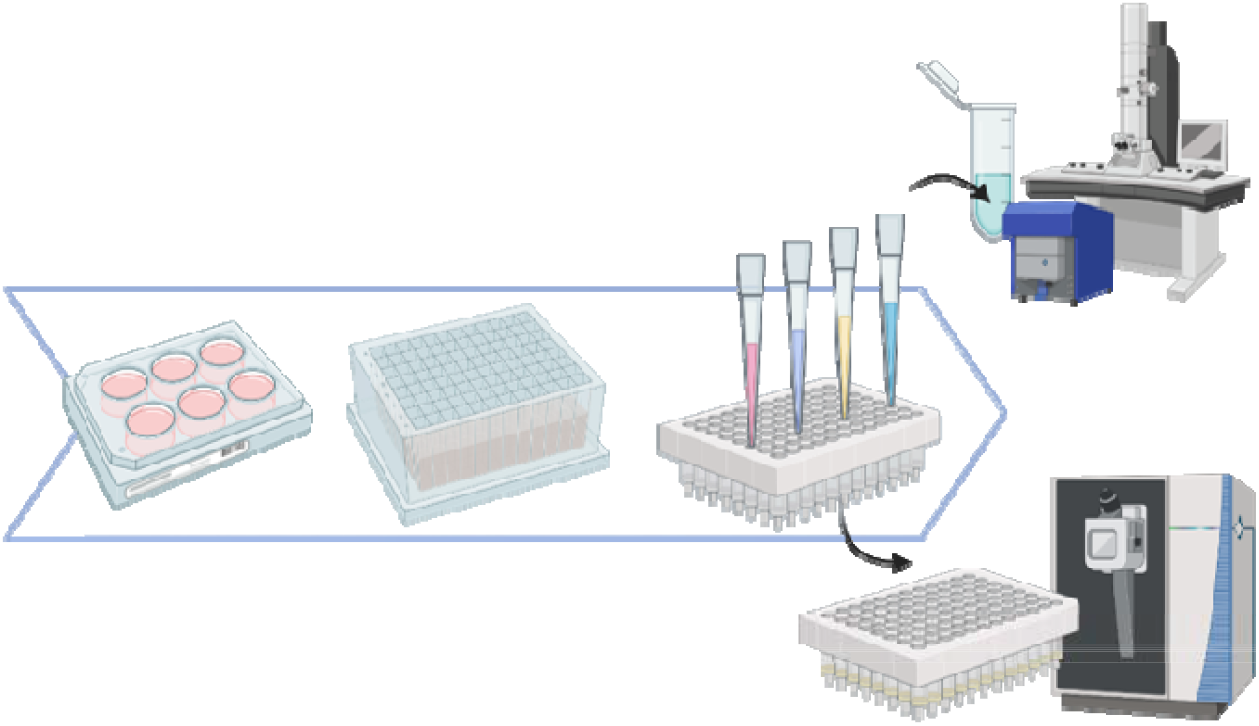

## 5. Introduction

According to the MISEV guidelines (1,2), extracellular vesicles (EVs) are defined as particles that are released by cells, are delimited by a lipid bilayer and cannot replicate on their own. EVs play crucial roles in intercellular communication, transfer of biomolecules, and modulation of biological processes. The EV umbrella term points to multiple subtypes of EVs that differ from one another by their biogenesis and size, including exosomes (30–150 nm) (3), microvesicles (100–1,000 nm) (4,5) and apoptotic bodies (500–2,000 nm) (6–8). Despite their differences, EV subtypes cannot be distinguished efficiently from one another due to the absence of specific markers or adequate separation techniques. Therefore, in scope of this study, we will limit the generic EV term to small EVs with a size (on average) below 200 nm. EVs carry a cargo of proteins, nucleic acids, lipids, and metabolites, reflecting the physiological or pathological state of their parental cells (9). As such, EVs in liquid biopsies (10) have emerged as promising diseases (stratification) biomarkers (11–13), to monitor or predict therapeutic responses (14,15) or as therapeutic agents (16,17). However, the protein complexity of the biofluid itself presents significant challenges for EV isolation and enrichment. Indeed, EVs in biofluids are typically present very high abundant non-EV-derived proteins such as albumin in plasma and serum, or as same-size particles such as lipoprotein particles (18,19). Therefore, for in-depth LC-MS/MS analysis of EV proteomes, robust EV enrichment strategies to achieve high purity and reproducibility are essential. Additionally, there is a growing interest in EV research towards scalable, high-throughput workflows for processing large sample numbers under various experimental conditions or for comparison of EVs isolated from different pathogenic cell lines

Ultracentrifugation (UC) is the most widely used EV isolation method according to the EV-TRACK public repository (20) as the method is simple, versatile and does not require specialized reagents, making it accessible to most labs. However, UC has limited throughput and is time-consuming, often requiring multiple consecutive rounds with extended centrifugation times. In addition, high centrifugal forces can cause EV aggregation, damage, or loss, potentially compromising downstream analyses. Furthermore, contamination with lipoproteins and protein aggregates remains a concern. Density gradient UC (dgUC) is often employed to improve purity by separating EVs from contaminants based on buoyant density (21). Despite its superior purity, dgUC is labor-intensive and has limited loading capacity and low throughput, although efforts have been made towards automation (22). Polyethylene glycol (PEG)-based precipitation reduces EV solubility, causing them to precipitate out of solution (23,24). PEG-kits have been made commercially available as they are popular for their simplicity, scalability, cost-effectiveness, and compatibility with a range of biofluids. However, such PEG-based methods often co-precipitate non-EV proteins and as they contain polymers, they may compromise the final EV purity and are also less suited for downstream LC-MS/MS analysis. Immunoaffinity capturing methods use antibodies targeting specific EV surface markers (e.g., CD9, CD63, or CD81) to capture EVs on magnetic beads or plates (25). Immunoaffinity offers high specificity, enriching EV subpopulations of interest (26) but then again may exclude EVs lacking the targeted markers, leading to biased enrichment (27). Although high-throughput applications are available, the availability and specificity of antibodies together with the overall cost remains a major concern (28). Size-exclusion chromatography (SEC) separates EVs based on their size using porous beads as a stationary phase from which EVs, being larger than most contaminants, elute earlier. SEC is gentle, preserving the integrity and functionality of EVs (29), although optimization of column dimensions and flow rates is necessary for inter-lab reproducibility (30,31). In addition, SEC has limited capacity, therefore requires either large column volumes for substantial sample input or reduction of the sample volume prior to EV enrichment, most often by ultrafiltration using 10 – 100 kDa MWCO filters (32). Indeed, ultrafiltration is most often used as a preparative concentration step prior to UC (33) dgUC (34) and SEC (32), but has not yet been established as a stand-alone EV enrichment strategy due to many challenges. These include reported poor EV purity, irreversible membrane blocking or fouling resulting in low passage of the biofluid (35). On the other hand, ultrafiltration is scalable, time-efficient, and suitable for high-throughput workflows.

Despite the existence of multiple EV enrichment strategies and hybrid approaches (33,36–38) accompanied with numerous comparative studies (39–42) no golden standard exists resulting in a case-by-case trade-off between scalability, cost and purity, yet also considering the available in-house expertise and equipment. Achieving high purity without compromising EV yield is a critical challenge for mass spectrometry-based proteomics where the presence of non-EV proteins can confound the proteomic depth due the stochastic competitive effect. Here, we evaluate a downscaled version of our filter-aided EV enrichment (FAEVEr) strategy (43) in a streamlined 96well format, including the pre-clearing of EV-containing samples, isolation and purification of the EVs as well as sample preparation for downstream LC-MS/MS analysis with minimal manual intervention. Our data show that intact EVs from conditioned medium are efficiently retained on the 300 kDa MWCO filter membrane and are successfully purified by including relatively high percentages of the TWEEN-20 detergent in the wash buffer, resulting in improved sensitivity and specificity starting from limited sample input.

## 6. Material and methods

### 6.1. Generation, isolation and pre-clearing of conditioned medium

#### 6.1.1. Recombinant EVs

The protocol for the transformation and production of recombinant extracellular vesicles (rEV) material is described in (44,45). In short, approximately 500,000 & 4 million HEK293T cells of low passage number (< 10) were seeded in a 6well or T75 falcon, respectively, with Dulbecco’s Modified Eagle Medium (DMEM, Gibco) containing 10% standard FBS and incubated for 48 h at 37 °C in 5% CO_2_. The HEK293T cells were transfected by adding a mixture of polyethyleneimine (PEI) and the DNA bait construct (12x, pMET7- GAG-eGFP, Addgene #80605) in DMEM + 2% FBS for 6 h at 37 °C in 5% CO_2_. The medium was then discarded and replaced with 1 mL (6well) or 8 mL (T75) fresh DMEM supplemented with 10% EV-depleted FBS (EDS, Thermo A2720801). After 48 h incubation at 37 °C in 5% CO_2_, cell transfection was evaluated under UV light (excitation at 488 nm and emission at 507 nm) before the conditioned medium (CM) was isolated.

#### 6.1.2. Pre-clearing conditioned media

For the comparison between the input, loading, wash and lysate fractions, 1 mL of CM from the 6well plate was pre-cleared individually by centrifugation at 1,000 x g for 5 min at room temperature (RT) and filtration using a 0.2 µm filtration 96well plate (Agilent, 204510-100) at 800 x g for 10 min, RT. The filtrate was collected in a 96well collection plate (Agilent, 5043-9308) and immediately used for EV enrichment by the FAEVEr 96well protocol (see below).

For the comparison of the EV proteomes using FAEVEr 96well and ultracentrifugation (UC) in combination with different percentages of TWEEN-20 in the wash buffer, 50 mL of GAG-eGFP transfected HEK293T cell CM was isolated and centrifuged at 1,000 x g for 15 min (RT). The supernatant was transferred to a syringe equipped with a 0.22 µm filter (Millex, SLGVR33RS) and gently pushed through. The filtrate was collected and divided per 600 µL to the corresponding wells (FAEVEr 96well) or UC tubes.

### 6.2 EV enrichment from pre-cleared conditioned medium

#### 6.2.1 FAEVEr 96well

600 µL of pre-cleared CM was loaded on a 300 kDa MWCO filter plate (Agilent, 201598-100). The plate was covered with an adhesive plastic foil, which was punctured to ensure consistent flow- through (FT), and mounted on a 96well collection plate. Loading the sample was performed in two steps as the loading capacity of the individual wells is limited to 350 – 400 µL. Both centrifugation steps were done at 1,000 x g for 10 min (RT). Once the complete sample was filtered, each well was washed with 200 µL 0.1% TWEEN-20 in PBS (pH 7.4), centrifuged at 1,000 x g for 7 min (RT). After three consecutive washing steps, the wells were washed once more with 200 µL 50 mM 4-(2- hydroxyethyl)-1-piperazineethanesulfonic acid (HEPES, pH 7.4) to remove residual washing buffer. Retained EVs were lysed by adding 50 µL lysis buffer (5% SDS in 50 mM triethylammonium bicarbonate (TEAB), pH 7.4) and the EV proteome recovered by centrifugation 1,000 x g for 5 min (RT) in a fresh 96well collection plate. The collected FT after loading and washing was either stored for proteome comparison, or discarded by inverting the collection plate. For EV characterization, filter-retained EVs were recovered by adding 200 µL 50 mM HEPES, shaking (1,000 rpm for 10 s) and vigorously pipetting up and down before transferring to a fresh Eppendorf.

For the optimization of the FAEVEr 96well protocol, different percentages of TWEEN-20 (0%, 0.1%, 0.5%, 1.0%, 2.5% and 5.0%) were used in the wash buffer to enrich and purify rEVs from 600 µL of rEV conditioned medium. Per TWEEN-20 concentration, eight replicates (48 in total) were prepared in parallel, from which two were used for characterization (NTA and TEM), one was used for Western blotting, three replicates were used for proteome analysis by LC-MS/MS, and the two remaining samples were stored.

#### 6.2.2 Ultracentrifugation

600 µL of pre-cleared CM was transferred to a 1 mL polycarbonate thick-wall tube and the volume was adjusted to 1 mL with PBS prior to ultracentrifugation at 100,000 x g for 90 min (4 °C). The supernatant was discarded manually and the pellet suspended in 1 mL washing buffer (PBS) before a second round of ultracentrifugation at 100,000 x g for 90 min (4 °C). The supernatant was again discarded manually. The EV enriched pellet was either suspended in 200 µL 50 mM HEPES for EV characterization or lysed with 50 µL lysis buffer and manually transferred to a fresh Eppendorf.

### 6.3 Proteome analyses

#### 6.3.1 Western blot

Samples for Western blot analysis were incubated for 5 min at 95 °C with XT sample buffer (4x) and XT reducing agent (20x) (BioRad). The protein material was separated by SDS-PAGE (4-12%) for 80 min at 120 V prior to transfer onto a PVDF membrane (100 V, 30 min). The membrane was incubated overnight at 4 °C with primary antibodies (rabbit monoclonal anti-Calnexin (ab133615) and mouse monoclonal anti-HIV p24 39/5 4A (ab9071); 1:1,000). The next day, the membrane was washed several times before incubation for 1 h at room temperature with secondary antibodies (goat anti-rabbit IRDye 680 CW and goat anti-mouse IRDye 800 CW; 1:10,000). Visualization was done on an Odyssey infrared imager (v3.0.16, LI-COR Biosciences).

#### 6.3.2. Sample preparation for LC-MS/MS analysis

All samples per experiment were prepared in parallel using the S-Trap 96well plate (Protifi, C02- 96well-10) following the manufacturer’s protocol. In short, the proteins were reduced using 5 mM tris(2-carboxyethyl)phosphine (TCEP) for 30 min at 55 °C and subsequently alkylated with 20 mM iodoacetamide (IAA) for 15 min at RT in the dark, before acidification with phosphoric acid (1.2% final concentration). The samples were diluted 1:7 with binding/wash buffer (90% methanol, 100 mM TEAB, pH 7.4), loaded on the S-Trap 96well plate and centrifuged for 5 min at 1,500 x g (RT). The trapped protein material was washed three times with 200 µL binding/wash buffer and centrifuged each time for 5 min at 1,500 x g (RT). 1 µg of trypsin per 125 µL 50 mM TEAB (pH 7.4) was added to the columns and digestion was performed overnight (approximately 16 h) at 37 °C after mounting the S-Trap plate to a fresh 96well collection plate. The next day, peptides were eluted in three consecutive centrifugation steps (1,500 x g for 5 min at RT) using 80 µL 50 mM TEAB (pH 7.4), 80 µL 0.2% formic acid in dH_2_O and 80 µL 2% ACN, 0.1% formic acid in dH_2_O. The eluates were transferred to MS vials and vacuum-dried in a SpeedVac vacuum concentrator.

#### 6.3.3. LC-MS/MS analysis

Peptides were re-dissolved in 25 µL loading solvent A (0.1% TFA in water/ACN (98:2, v/v)) and the peptide concentration was determined on a Lunatic instrument (Unchained Lab). For each sample the injection volume was adjusted to inject equal amounts of peptide material (500 ng) or maximized (15 µL) when less material was available. The peptide material was injected for LC-MS/MS analysis on a Vanquish™ Neo UHPLC System in-line connected to an Orbitrap Exploris 240 mass spectrometer (Thermo). Injection was performed in trap-and-elute workflow in combined control mode (maximum flow of 60 µl/min and maximum pressure of 800 bar) in weak wash solvent (WW, 0.1% trifluoroacetic acid in water/acetonitrile (ACN) (99.5:0.5, v/v)) on a 5 mm trapping column (Thermo scientific, 300 μm internal diameter (I.D.), 5 μm beads). The peptides were separated on a 250 mm Aurora Ultimate, 1.7µm C18, 75 µm inner diameter (Ionopticks) kept at a constant temperature of 45 °C. Peptides were eluted by a gradient starting at 0.5% MS strong wash solvent (SW) (0.1% formic acid (FA) in acetonitrile) reaching 26% MS SW in 30 min, 44% MS SW in 38 min, 56% MS SW in 40 min followed by 5-minute wash at 56% MS SW and column equilibration in pressure control mode (separation column: fast equilibration, maximum pressure of 1,500 bar, equilibration factor of 2; trap column: fast wash and equilibration, wash factor=100) with MS WW. The flow rate was set to 300 nl/min. The mass spectrometer was operated in data-independent mode, automatically switching between MS and MS/MS acquisition. Full-scan MS spectra ranging from 400-900 m/z with a normalized target value of 300%, a maximum fill time of 25 ms and a resolution at of 60,000 were followed by 30 quadrupole isolations with a precursor isolation width of 10 m/z for HCD fragmentation at an NCE of 30% after filling the trap at a normalized target value of 2000% for maximum injection time of 45 ms. MS2 spectra were acquired at a resolution of 15,000 with a scan range of 200-1,800 m/z in the Orbitrap analyser without multiplexing. The isolation intervals were set from 400 – 900 m/z with a width of 10 m/z using window placement optimization. EASY-ICTM was used in the start of the run as Internal Mass Calibration.

#### 6.3.4. Database search

Data-independent acquisition (DIA) spectra were searched with the DIA-NN software (v1.9.2) (46) in library-free mode against the combination of two protein databases downloaded from Swiss- Prot; the complete human protein sequence database (January 2021, 20,394 sequences) supplemented with bovine serum protein sequences (March 2021, 618 sequences). For searches that included rEV material, we included the GAG-eGFP protein sequence as well. The mass accuracy was set to 10 ppm and 20 ppm for MS1 and MS2, respectively, with a precursor FDR of 0.01. Enzyme specificity was set to trypsin/P with a maximum of two missed cleavages. Variable modifications were set to oxidation of methionine residues (to sulfoxides) and acetylation of protein N-termini. Carbamidomethylation of cysteines was set as a fixed modification. The peptide length range was set to 7-30 residues with a precursor charge state between 1 and 4, and an m/z range between 400-900 and 200-1,800 for the precursor and fragment ions, respectively. Cross-run normalization was set to RT dependent with the quantification strategy set to high accuracy and the neural network classifier to single-pass mode. Matching between runs was allowed for the comparison of the washing buffers and the melanoma EVs, but not for the comparison of the FT fractions. The result file was further processed in KNIME (v4.3.3) by removing non-proteotypic peptides and protein identifications with a q-value above 0.01 or with less than two precursor identifications. Peptide quantifications were aggregated to protein group quantifications using the median of the corresponding normalized LFQ values. Further data analysis was performed with Perseus (version 1.6.14.0) (47), GraphPad Prism (v9.4.1) and RStudio (2023.12.1). For iBAQ quantifications, the publicly available DIA-GUI was used with default settings (48).

### 6.4 Data analysis

#### 6.4.1. Comparison of relative and absolute protein abundances

The absolute protein abundance (iBAQ) values were plotted using the average log_2_ intensities per experimental setup for proteins with at least three valid values in the corresponding group. For the comparison of the absolute and relative intensities of the total ion chromatogram, the raw intensities from human and bovine precursors were selected and plotted. For the number of protein identifications and constructing the Venn diagram, the proteins that were represented in each replicate per fraction were plotted. For the EV marker abundance, LFQ intensities were used. For the multiple sample comparison, proteins from the individual fractions were filtered for at least three valid values in at least one group prior to imputation of the missing values (width 0.3; down shift 1.8). Significantly differentially abundant proteins (p-value < 0.001) were selected and hierarchically clustered (Euclidean distance with averaged linkage) after Z-scoring. Protein clusters were manually distinguished and selected for gene ontology analysis using the online Webgestalt tool (49) (no redundant terms, FDR 0.05) or STRING database. Protein networks of interest were processed in Cytoscape (v3.10.1). For two-sample comparisons, the corresponding groups were selected and proteins with at least three valid values in minimum one group were selected prior to imputation of the missing values (width 0.3; down shift 2.2). Proteins with a -log_10_(p-value) > 1.3 and fold change > 2 (log_2_ difference > |1|) were considered to be significantly differentially abundant.

### 6.5. EV characterization

#### 6.5.1. Nanoparticle tracking analysis

Recuperated EV particles were diluted to 1 mL using PBS (pH 7), injected in a calibrated Zetaview and analyzed with the corresponding software package (version 8.05.16 SP2). The temperature was maintained at 23 °C and the analysis was done at a sensitivity of 70 and a shutter of 100 for 3 cycles at 60 fpm. The results were plotted and annotated in GraphPad Prism (v10.2.2)

#### 6.5.2. Transmission electron microscopy

The recovered EV fractions were concentrated in a SpeedVac vacuum concentrator to approximately 15 – 20 µL. Aliquots (5 µl) of the EV solutions were blotted for 1 min on formvar- and carbon-coated Ni maze grids (EMS), which were glow discharged for 40 s at 15 mA. The grids were then washed five times in droplets of dH_2_O and stained in a droplet of 1/4 Uranyl Acetate Replacement Stain (EMS)/ dH_2_0 for 1 min. Excess stain was removed with filter paper and the grids were air dried for at least four hours before viewing with the TEM. Imaging was done at 80 kV on a JEM1400plus (JEOL).

## 7. Results

### 7.1. Optimization of the high-throughput EV enrichment strategy

#### 7.1.1. Workflow description and considerations

Our filter-aided extracellular vesicle enrichment (FAEVEr) strategy on a complete 96well format is illustrated in **FIGURE 1**. In short, conditioned medium is pre-cleared by centrifugation and filtration (0.2 µm pore size). The filtrate is collected in 96well collection plate and either frozen (−80 °C) or processed immediately. For EV enrichment, the pre-cleared conditioned medium is loaded on a 300 kDa MWCO 96well filter plate and centrifuged. The retentate is then washed three times to remove contaminating non-EV protein material. Purified particles are recovered from the filter membrane for characterization by NTA and TEM. For protein analysis, the EVs are immediately lysed on the filter membrane using 5% SDS. Upon the retrieval of the EV proteome by centrifugation in a 96well collection plate, the individual samples are prepared in parallel following the S-Trap 96well protocol, thus maintaining the streamlined high-throughput approach.

**Figure 1:**
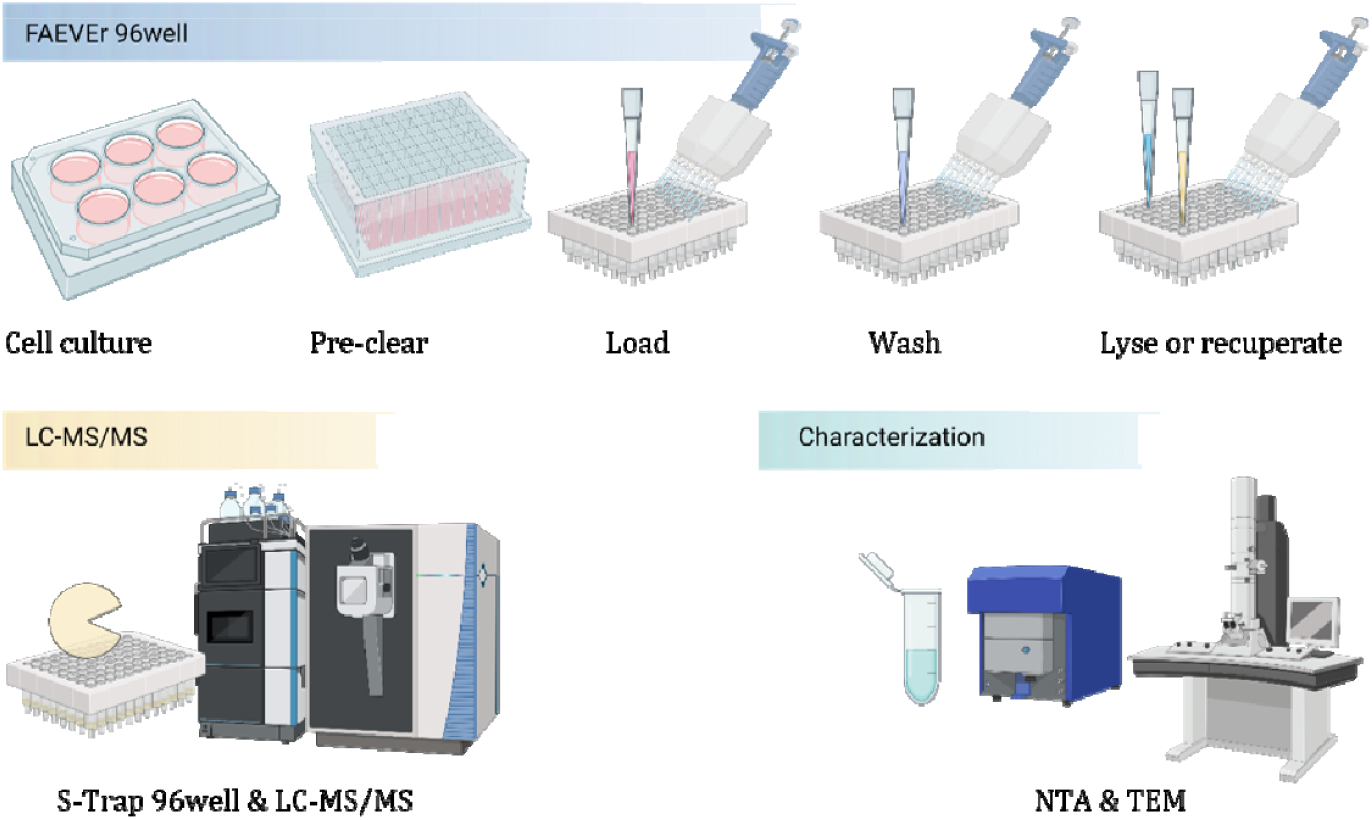
Description of the FAEVEr 96well workflow. Conditioned medium containing 10% EV-depleted FBS is isolated and pre-cleared by centrifugation and filtration (0.2 µm filter) in a 96well format. The filtrate is collected and either frozen (−80 °C) or processed immediately. After loading the pre-cleared conditioned medium on a 300 kDa MWCO 96well filtration plate, the plate is centrifuged. The retained particles are purified by three consecutive wash steps. For LC-MS/MS analysis, the purified EVs are lysed on the filter (yellow) followed by the collection of the proteome by centrifugation and processing by S-Trap 96well. For characterization, the particles are recuperated from the filter (cyan) after the wash steps and analyzed by NTA and TEM.

A major, and justified, concern of dead-end ultrafiltration (UF) is the irreversible fouling or clogging of the membrane due to accumulation of protein material on the filter membrane. This results in a poor and a non-consistent flow-through, or even sample loss. We found this to be even more pronounced using a 96well format with a filter surface of 0.2 cm^2^ as smaller surfaces are more susceptible to fouling, making consistent filtration a major concern. Despite 100 kDa MWCO filters being most prominently used within the EV field as a concentration step during SEC or dgUC mediated EV enrichment, UF has not yet acquired foothold as a stand-alone enrichment strategy. We argued that 300 kDa MWCO filters, with a pore size of roughly 30 – 35 nm according to the manufacturer, were more applicable for a consistent EV enrichment yielding EVs with sufficiently high purity. Indeed, with an average size range between 50 and 150 nm, exosomes should be quantitatively retained, whereas nearly all globular (human) proteins would not **(FIGURE S1)**. This in contrast to 100 kDa MWCO, where approximately 13% of the human proteome would accumulate on the membrane. To our knowledge, 300 kDa MWCO membrane are only available in polyethylene sulfone (PES) material, which comes with the additional benefits of being hydrophilic with reduced non-specific protein adsorption, resulting in a high flow rate. In addition, PES is resistant to a wide array of chemicals (e.g. detergents) and a pH range of 1 – 14 (according to the manufacturer).

#### 7.1.2. Extracellular vesicles are successfully retained on the filter membrane

To validate that our approach on a 300 kDa MWCO 96well filter plate was capable of quantitatively retaining intact EV particles, we used recombinant EVs (rEVs) as a reference source (44,50). We initially compared the proteomes of the non-enriched crude conditioned medium with three different fractions that were collected after filtration; the flow through after sample loading, the washing and the lysate steps. Here, an exploratory 0.1% TWEEN-20 was added to the wash buffer. We specifically focused on the abundance of the general rEV markers (including luminal Gag-eGFP) as well as differentially abundant or uniquely identified proteins between the individual fractions. In addition, as rEVs were harvested from conditioned medium containing 10% EV depleted FBS (EDS), we considered bovine proteins as a direct measure of contamination. Retained rEVs were characterized by transmission electron microscopy (TEM), nanoparticle tracking analysis (NTA) and Western blotting for GAG-eGFP **(FIGURE 2 A-C)** and the proteomes were analyzed by LC-MS/MS (**FIGURE S2**).

**Figure 2:**
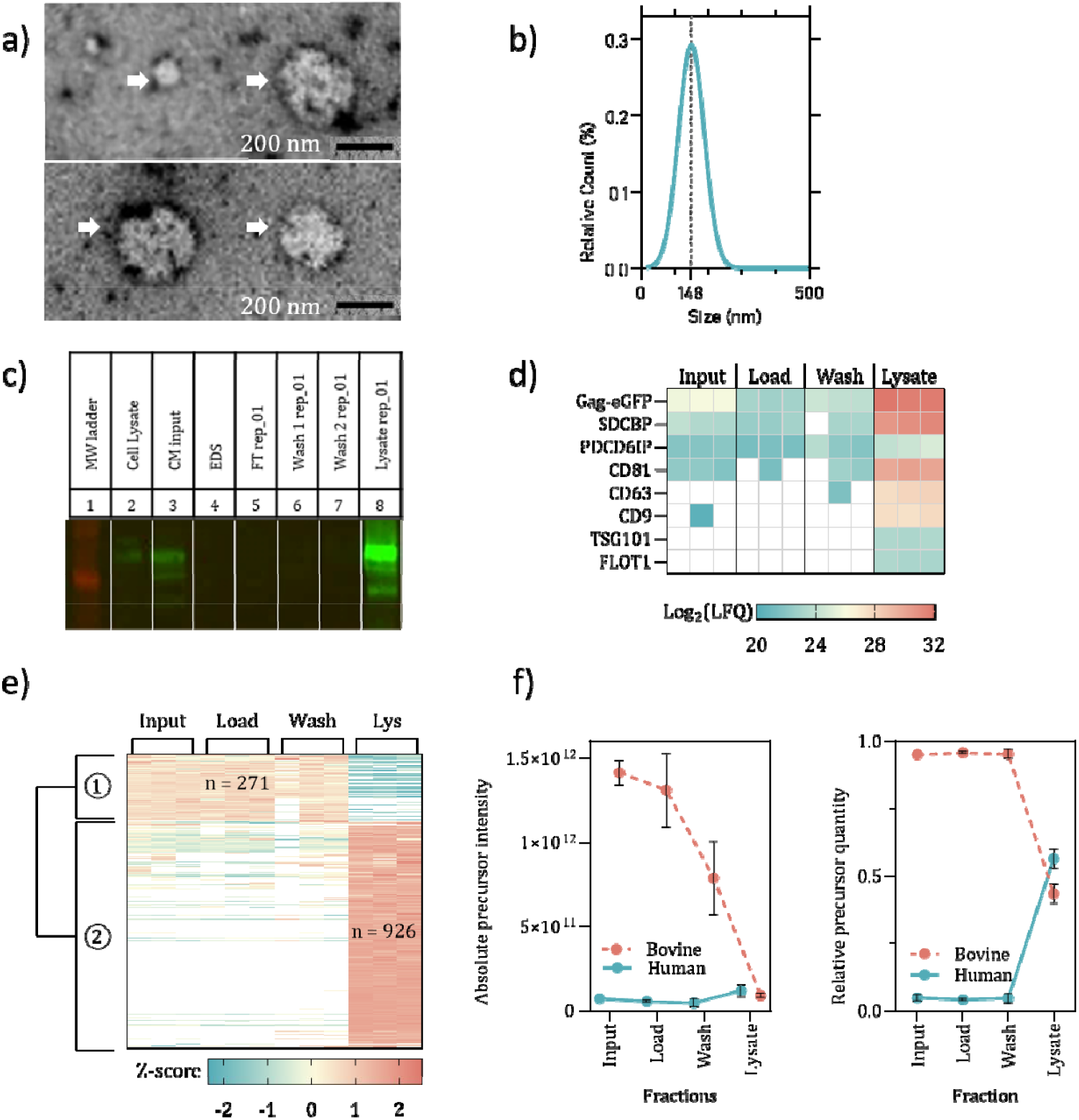
Recombinant EVs are successfully enriched and purified by FAEVEr using a 300 kDa MWCO 96well filter plate. rEV particles were enriched and recuperated from the filter for characterization by (a) TEM and (b) NTA. (c) Western blot analysis of the individual fractions shows that no luminal overexpressed Gag- eGFP is observed in the initial flow-through or wash, but is highly abundant in the enriched rEVs. (d) Eight rEV protein markers were searched across the different fractions after LC-MS/MS analysis, indicating a high abundance (Log2(LFQ) intensities) in the lysate fraction. (e) Multiple– sample testing between the individual fractions revealed two major clusters of proteins that were more abundant in the lysate fraction compared to the other fractions. (f) Absolute and relative Precursor intensities originate largely (>96%) from bovine precursors in the input, load and wash fractions, but are greatly reduced in the lysate fraction (43%).

Upon proteome analysis, all eight rEV-specific molecular protein markers (GAG-eGFP, SDCBP, PDCD6IP, CD81, CD63, CD9, TSG101 and FLOT1) were found to be highly abundant in the lysate, whereas they were generally low abundant or completely devoid in the other fractions **(FIGURE 2D)**. Differences in protein abundance between the fractions was assessed by multiple sample testing followed by hierarchical clustering of the significantly differentially abundant proteins (p- value < 0.01) **(FIGURE 2E)**. We found that 926 proteins were significantly higher abundant in the lysate fraction, which included over 550 unique proteins and are predominantly associated with EV biogenesis, including MVB sorting, ESCRT and SNARE complex and small GTPase mediated signal transduction **(FIGURE S3)**. In contrast, 271 proteins, of which 93 were bovine proteins, were significantly less abundant or depleted all together from the lysate fraction, with human proteins being mainly associated with collagen synthesis, extracellular matrix constituents/organization and carbohydrate metabolism. Concerning contaminating bovine proteins, we found that the input, loading and wash fraction were dominated by spectra originating from bovine precursors (> 96% of the total ion chromatogram, TIC), which was greatly reduced in the lysate fraction (app. 43% of the TIC) **(FIGURE 2F)**.

### 7.2. 5% TWEEN-20 in the wash buffer is optimal for the identification of EV associated proteins

After validating that FAEVEr 96well was capable of retaining intact EVs while removing the bulk of non-EV proteins, we assessed the optimal concentration of TWEEN-20 in terms of overall EV purity. In addition, through the parallel enrichment of 48 samples within a limited timeframe (about 2 h) using minimal resources, we showcase the high-throughput potential of the 96well FAEVEr approach. After enriching rEVs from 600 µL conditioned medium each followed by purification using 0%, 0.1%, 0.5%, 1.0%, 2.5% and 5% TWEEN-20 in the wash buffer in octuplets, we recuperated particles in duplicate for characterization (NTA and TEM), and recovered the proteome in triplicate for LC-MS/MS analysis. Two remaining samples were stored for future reference. In parallel, we enriched rEV particles using ultracentrifugation using the same starting volume.

Particle count and size distribution were determined by NTA revealing a typical size distribution for sEV particles, ranging between 145 and 160 nm on average (**FIGURE 3A and FIGURE S4)**. TEM imaging showed the distinct morphology of EVs, thereby validating that EVs remained intact on the filter during their enrichment (**FIGURE 3B**). Incomplete recovery of EVs from the filter membrane is a known caveat of ultrafiltration both during EV enrichment or concentration prior to, for instance, SEC and dgUC (35). However, for LC-MS/MS analysis it is unnecessary to recuperate the intact EVs back from the filter as we recover the EV proteome through immediate on-filter lysis using high percentages of SDS followed centrifugal elution

**Figure 3:**
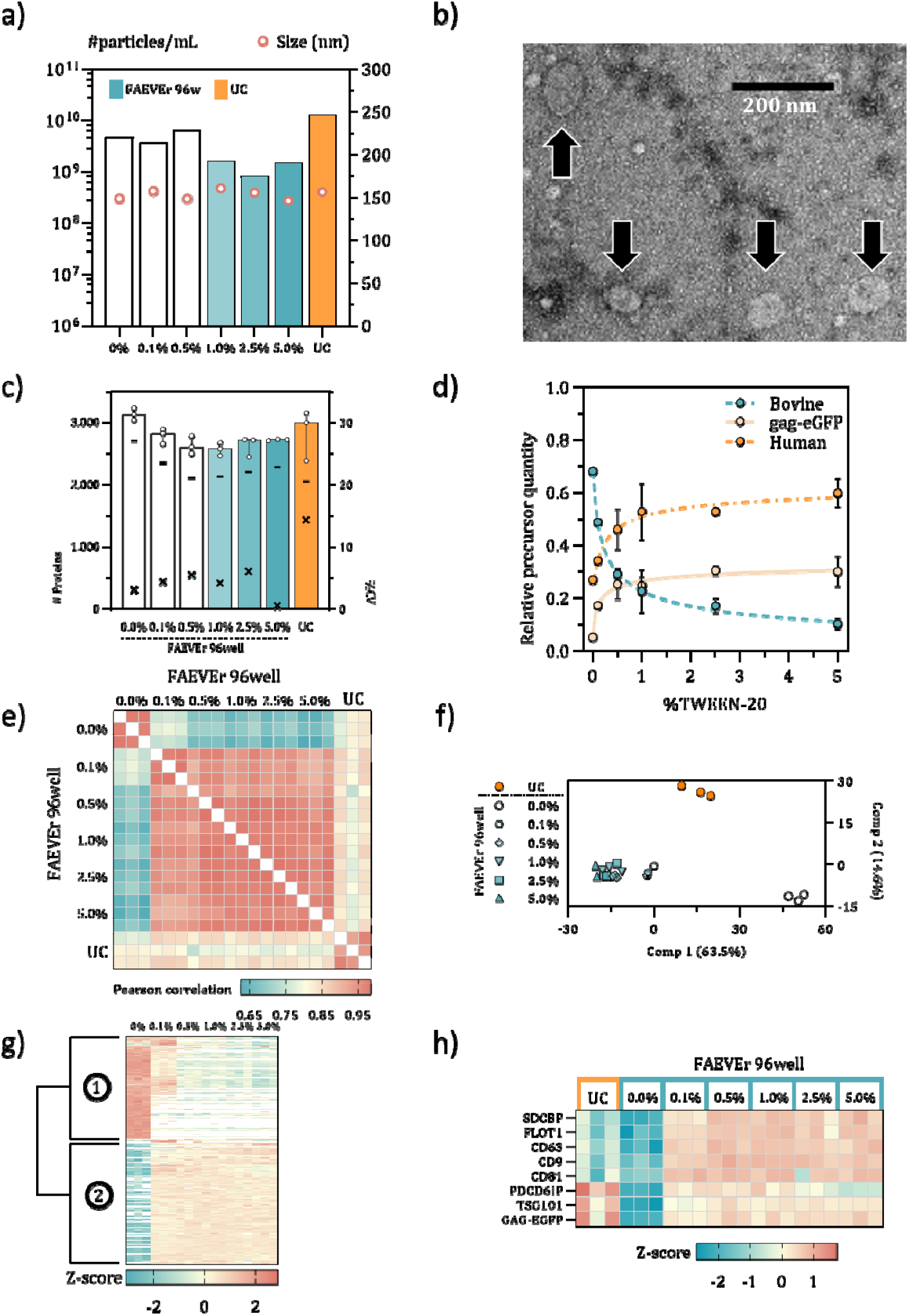
Comparison of the effect of the TWEEN-20 concentration on the rEV purity. Retained rEVs were characterized by NTA (a) and TEM (b). (c) Number of protein identifications (spheres) with the number of proteins identified across all replicates (n = 3) indicated by the dash. The corresponding coefficient of variance (%CV) is indicated by ‘x’ on the right y-axis. (d) Bovine precursor abundance decreases exponentially proportional to the concentration of TWEEN-20; from nearly 70% at 0% TWEEN-20 to 10% using 5% TWEEN-20 in the washing buffer. (e) High Pearson correlation between the different replicates and clustered grouping of enrichment strategies (f) suggests high reproducibility. (g) Hierarchical clustering of significantly differentially abundant proteins reveals two major clusters. (h) Eight protein markers specific for rEVs were found across all the experiments with the transmembrane (CD63, CD9 and CD81), peripheral (FLOT1) and membrane-associated (SDCBP) proteins being relatively higher abundant in the FAEVEr with TWEEN-20 compared to the luminal markers (PDCD6IP, TSG101 and Gag-eGFP) in ultracentrifugation. FAEVEr 96well without TWEEN-20 has the overall lowest relative abundance of rEV markers.

### 7.3. EV markers but not contaminants dominate the proteome in FAEVEr 96well with TWEEN-20

The obtained LC-MS/MS data were analyzed for specificity, sensitivity and robustness to determine which TWEEN-20 concentration in the washing steps is optimal for downstream proteome analysis of the retained EVs. First, the number of protein identifications was plotted, with differentiation between human and bovine proteins. The highest number of protein identifications is observed in FAEVEr 96well without TWEEN-20 in the wash buffer, but immediately declines when TWEEN-20 is added, even at very low (0.1%) concentrations and plateaus towards higher percentages of TWEEN-20 **(FIGURE 3C and S6a)**.. Furthermore, the %CV appears to be low (≤ 6%) for all FAEVEr experiments. In comparison, the ultracentrifugation approach reaches a similar number of protein identifications (appr. 3,000), from which only 65% is identified across all replicates with a %CV of 14.31%. Although the absolute number of bovine protein identifications does not dramatically decreases (**FIGURE S6b)**, we investigated their abundance in terms of precursor intensities. We observed an exponential decrease in bovine precursor abundance proportional to the concentration TWEEN-20, from nearly 70% at 0% TWEEN-20 to just 10% using 5% TWEEN-20 in the washing buffer **(FIGURE 3D)**. As an illustration, the absolute abundance (iBAQ intensity) of contaminating bovine apolipoproteins decreases with an increasing percentage of TWEEN-20 (**FIGURE S5**). To assess the technical reproducibility of the approach, we investigated the overall correlation between the individual samples **(FIGURE 3E)** and performed a principal component analysis (n = 1,163, **FIGURE 3F**). Both plots revealed a high reproducibility between the replicates with an average Pearson correlation between 0.94 and 0.98 and condensed clustering on the PCA plot. We noticed a large variability between three major groups: FAEVEr in the absence and presence of TWEEN-20, and ultracentrifugation.

To investigate the overall protein heterogeneity and protein levels in FAEVEr-prepared samples with or without TWEEN-20, a multiple sample test was performed from which the significantly differential proteins (p-value < 0.001) were isolated, z-scored and hierarchical clustered, resulting in two main clusters (**FIGURE 3G**). Cluster 1 contains 693 proteins that are more abundant in the 0% TWEEN-20 washes, with 109 being of bovine origin. Gene ontology (GO) analysis of the human proteins revealed significant association with the extracellular matrix (e.g. collagen and fibril), mRNA processing (e.g. spliceosome), protein folding (e.g. chaperone activity) and metabolic processes. Note that a large part of the proteins is (nearly) completely absent in samples prepared with higher (>0.1%) percentages of TWEEN-20. Cluster 2 includes GO terms such as MVB sorting pathway, vesicle transport/fusion and GTPase activity are highly enriched. Interestingly, out of the 825 proteins, 317 are TM proteins. The latter is of specific interest as endocytosis, the shuttling of TM plasma membrane proteins to the endosome, is closely intertwined with EV biogenesis. In addition, TM proteins are generally more challenging to identify by LC-MS/MS, although they are of specific interest for drug development and diagnostics.

Specifically, FAEVEr 96well with 5% TWEEN-20 appeared to be very performant with a consistent high number of identifications between the replicates with no missing values (2,272 out of 2,842 proteins, 80%), low %CV (<2%) and high inter-sample correlation (97%). In addition, this setup has the least interference of bovine precursors (10%) and non-EV proteins resulting in a high abundance of true EV associated proteins such as the ESCRT machinery, transmembrane proteins and the commonly used EV protein markers (**FIGURE 3H**).

### 7.4. Comparison FAEVEr 96well with ultracentrifugation

To further examine the potential of FAEVEr 96well for EV enrichment towards proteomics, we compared our FAEVEr 96well approach in combination with 5% TWEEN-20 with differential ultracentrifugation (UC) using an equal starting volume. To highlight the differences in relative protein abundances between the canonical UC enrichment and FAEVEr 5% TWEEN-20 on a 96well format, we performed a two-sample comparison of 2,775 proteins after filtering for at least three valid values in minimum one group (**FIGURE 4**). 1,049 proteins were significantly differentially abundant (p-value < 0.05 and log2 > |1|), with 862 and 225 proteins being higher abundant (or uniquely identified) in UC and FAEVEr, respectively. In UC, 112 (14%) and 106 (13%) are secreted proteins from bovine and human origin, respectively. In contrast, in FAEVEr with 5% TWEEN-20 only 2 bovine and 17 human secreted proteins were found, suggesting an improved removal of contaminating proteins. Interestingly, only 64 proteins in UC (8%) were TM proteins compared to 123 (56%) in FAEVEr. GO analysis revealed that in UC these TM proteins were specifically localized within the Golgi and ER membrane, compared to FAEVEr 5%, in which TM proteins were specifically associated with TM transport, adhesion and localization **(FIGURE S7a & S7b)**. Finally, intracellular proteins were present in the UC approach (519 or 63%) to a far larger extent compared to FAEVEr (72 or 30%). In UC, the intracellular proteins were associated with DNA replication and cytoplasmic translation. Moreover, 137 of these intracellular proteins were uniquely identified in UC and grouped specifically around RNA processing, the spliceosome and the nucleus. In FAEVEr, however, the intracellular proteins were predominantly associated with the ESCRT machinery and MVB assembly.

**Figure 4:**
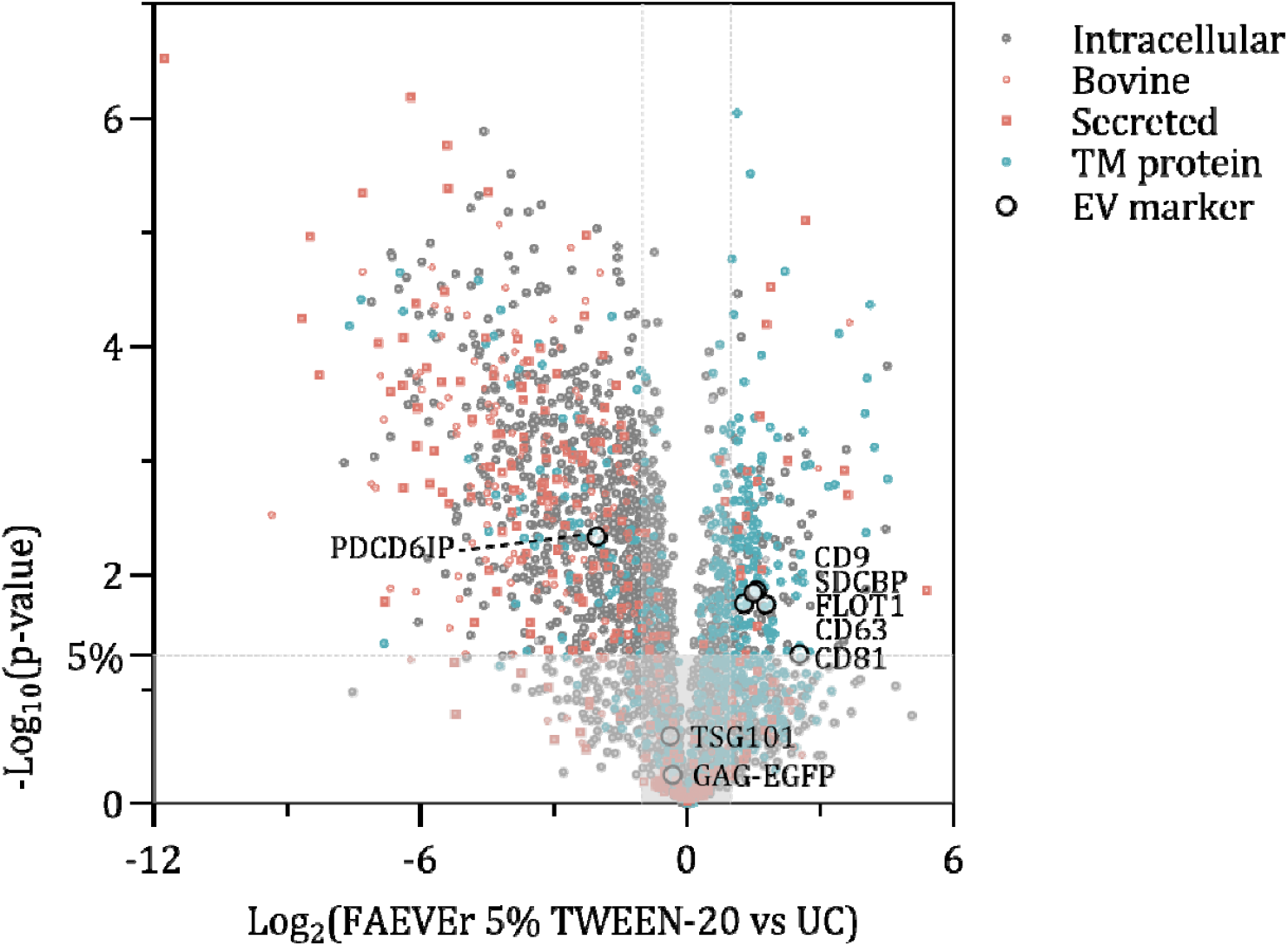
Quantitative comparison of the identified proteins between FAEVEr 96well in combination with 5% TWEEN-20 versus EVs isolated by ultracentrifugation. Volcano plot of the log_10_(p-values) and corresponding log_2_ differences indicate that in FAEVEr 96well with 5% TWEEN-20, the transmembrane protein were relatively more abundant whereas bovine and human secreted proteins were significantly decreased, compared to the proteomes of EVs isolated by ultracentrifugation.

## 8. Discussion

Here, we established the downscaling of the FAEVEr methodology using 300 kDa MWCO filters to a 96well format towards high-throughput enrichment of EV particles from conditioned cell medium for mass spectrometry-based proteomics. Due to the limited sample input and small filtration membrane surface, we re-evaluated the purity of recovered EV lysate after implementing TWEEN- 20 in the wash buffer. We concluded that EV enrichment from a mere 600 µL starting material successfully retains intact EVs as illustrated by the characterization through size distribution (by NTA) and morphology (by TEM). In addition, we provide insights in the efficient recovery of the EV proteome by immediately lysing the retained EVs on the filter resulting in a high abundance of specific EV markers, EV biogenesis associated proteins and transmembrane proteins through LC-MS/MS analysis. In addition, through quantitative comparison between the input, loading, wash and lysate fraction, we conclude that no or little EV specific material was lost during subsequent round of centrifugation whereas a large part of the bovine serum proteins is removed when adding a trace (0.1%) of TWEEN-20 to the wash buffer. Of note, this experiment was conducted with 1 mL of conditioned medium per replicate, derived from 6well plates, from which 600 µL was used for EV enrichment. This highlights the throughput advantage over other EV enrichment techniques, such as density gradient ultracentrifugation, which often require large starting volumes of conditioned medium, which can hamper particularly when incorporating multiple replicates or experimental conditions.

By comparing the EV proteome after completing the FAEVEr 96well protocol using different percentages of TWEEN-20 in the washing buffer, we confirm that ultrafiltration without additives in the washing buffer (0% TWEEN-20 in PBS) results in poor purity, yet high recovery, based on the NTA and proteomics results. This is in line with previous observations where membrane fouling was described as a major drawback of ultrafiltration, resulting in impure EV proteomes. We established that including TWEEN-20 in the wash buffer has a profound impact on the sample purity and is directly proportional to the final concentration TWEEN-20 (**FIGURE 2C**). As dead-end ultrafiltration retains all material with a size surpassing the 300 kDa MWCO threshold, it is susceptible to co-isolation of lipoprotein particles. By comparing the corresponding absolute abundance (iBAQ) of apolipoproteins as a measure of apolipoprotein particle abundance across the different experiments, we observed a decreasing trend with increasing percentages of TWEEN-20 in FAEVEr 96well, even reaching levels well-below the UC approach (**FIGURE S5**). However, it appears that near-complete depletion of lipoprotein particles from the EV sample requires more specific, yet tedious, two-dimensional approaches, e.g. SEC in combination with density gradient UC (22) or filtration in combination with electrophoretic separation (51). Again, a case-by-case trade-off has to be considered.

We established from our comparison that including 5% TWEEN-20 is optimal to minimize non-EV protein interference, including bovine serum proteins. This results in an increased relative abundance of proteins related to EV biogenesis and membrane proteins. Surface exposed membrane proteins, and in particular TM proteins, are highlighted in the EV biomarker field as they reflect the surface of the parental (diseased) cell (52) and have the potential of being easily picked up in liquid biopsies using antibodies. In addition, TM proteins are expected to be highly prevalent in EVs compared to luminal proteins. Indeed, EVs have an exceptional high relative surface to volume ratio and the EV biogenesis is closely intertwined with endocytosis, the constitutive transport of plasma membrane TM proteins to the endosome. In addition, intact TM proteins, containing at least one hydrophobic TM helix, are very unlikely to be present as such in conditioned media without a lipid bilayer for embedment as they would aggregate, precipitate and therefore be removed during the protocol.

We compared the FAEVEr 96well methodology, including 5% TWEEN-20 washes, with the widely implemented UC approach, and observed not only a depletion of bovine serum and human secreted proteins, but also of intracellular proteins associated with RNA processing (in particular the spliceosome), chaperone mediated protein folding and mitosis (DNA replication). We hypothesize that these particular protein groups might in fact not reside within the EV lumen, but rather largely originate from unavoidable cell death during cell culture, and are thereby subsequently co-enriched due to incomplete removal and therefore wrongly considered to be EV proteins, which will bias data interpretation. However, further research is required to validate this presumption.

## 9. Conclusion

The FAEVEr 96-well strategy for enriching EVs from conditioned medium (CM) has proven highly effective in retaining intact EVs while enabling the quantitative removal of non-EV proteins by incorporating TWEEN-20 in the wash buffer. A systematic comparison using an established reference EV material of increasing TWEEN-20 concentrations revealed optimal purity and reproducibility at 5% TWEEN-20. Although previous studies by Osteikoetxea *et al*. (53) demonstrated that small EVs remain intact in buffers containing 10% TWEEN-20, we found that concentrations >5% adversely affected flow-through due to increased viscosity. Our results demonstrate that abundant xenoproteins (bovine serum) and non-EV proteins (secreted proteins) are quantitatively removed, thereby enhancing the sensitivity and specificity to identify less abundant, but biologically relevant proteins, such as transmembrane and EV biogenesis-related proteins. Moreover, the FAEVEr 96-well platform exhibited exceptional robustness and scalability, processing 36 and 84 samples in parallel during the TWEEN-20 comparison and melanoma study, respectively, within a short timeframe and resulted in very high reproducibility. The complete EV enrichment workflow (including the isolation of CM followed by pre-clearing, EV enrichment, and purification) was executed in approximately 3 hours under low centrifugation speeds (up to 1,200 × g) using standard lab equipment. By integrating the commercial S-Trap 96-well plates for EV proteome preparation, we also achieved high protein recovery using elevated SDS percentages, while further streamlining the workflow in a high-throughput 96-well format. This setup is ideally suited for large-scale EV proteome screening from cell culture, ensuring high reproducibility without having to compromise between replicates and experimental conditions.

## 4. List of abbreviations

CM: Conditioned medium
dgUC: Density gradient ultracentrifugation
DMEM: Dulbecco’s Modified Eagle Medium
EDS: EV-depleted foetal bovine serum
EV: Extracellular vesicle
FAEVEr: Filter-aided extracellular vesicle enrichment
FBS: Foetal bovine serum
FT: Flow-through
HEPES: 4-(2-hydroxyethyl)-1-piperazineethanesulfonic acid
LC-MS/MS: Liquid chromatography coupled tandem mass spectrometry
MISEV: Minimal information for studies of extracellular vesicles
MWCO: Molecular weight cut-off
NTA: Nanoparticle tracking analysis
PEG: Polyethylene glycol
PEI: Polyethyleneimine
PES: Polyethylene sulfone
rEV: Recombinant extracellular vesicles
RT: Room temperature
SDS: Sodium dodecyl sulphate
SEC: Size-exclusion chromatography
TEAB: Triethylammonium bicarbonate
TEM: Transmission electron microscopy
UC: Ultracentrifugation

## 14. Author contributions

JP planned, designed and performed the experiments and wrote the manuscript. FDM and DDP assisted with the experiments. TVDS performed the HEK293T cell culture and production of rEV material. FB performed the electron microscopy. All authors read and approved the manuscript.

15. Acknowledgements

The authors thank all operators of the VIB Proteomics Core for their excellence and input. The authors thank the lab of An Hendrix for kindly providing us with the required purified rEV aliquots.

## 16. Data availability

The mass spectrometry proteomics data have been deposited to the ProteomeXchange Consortium via the PRIDE partner repository with the dataset identifiers PXD059819 (optimization FAEVEr 96well). We have submitted all relevant data of our experiments to the EV-TRACK knowledgebase (EV-TRACK ID: EV250006) (20).

