## Supplementary material for "A 96well ultrafiltration approach for the high-throughput proteome analysis of extracellular vesicles isolated from conditioned medium": Suppl figures

### Slide 1
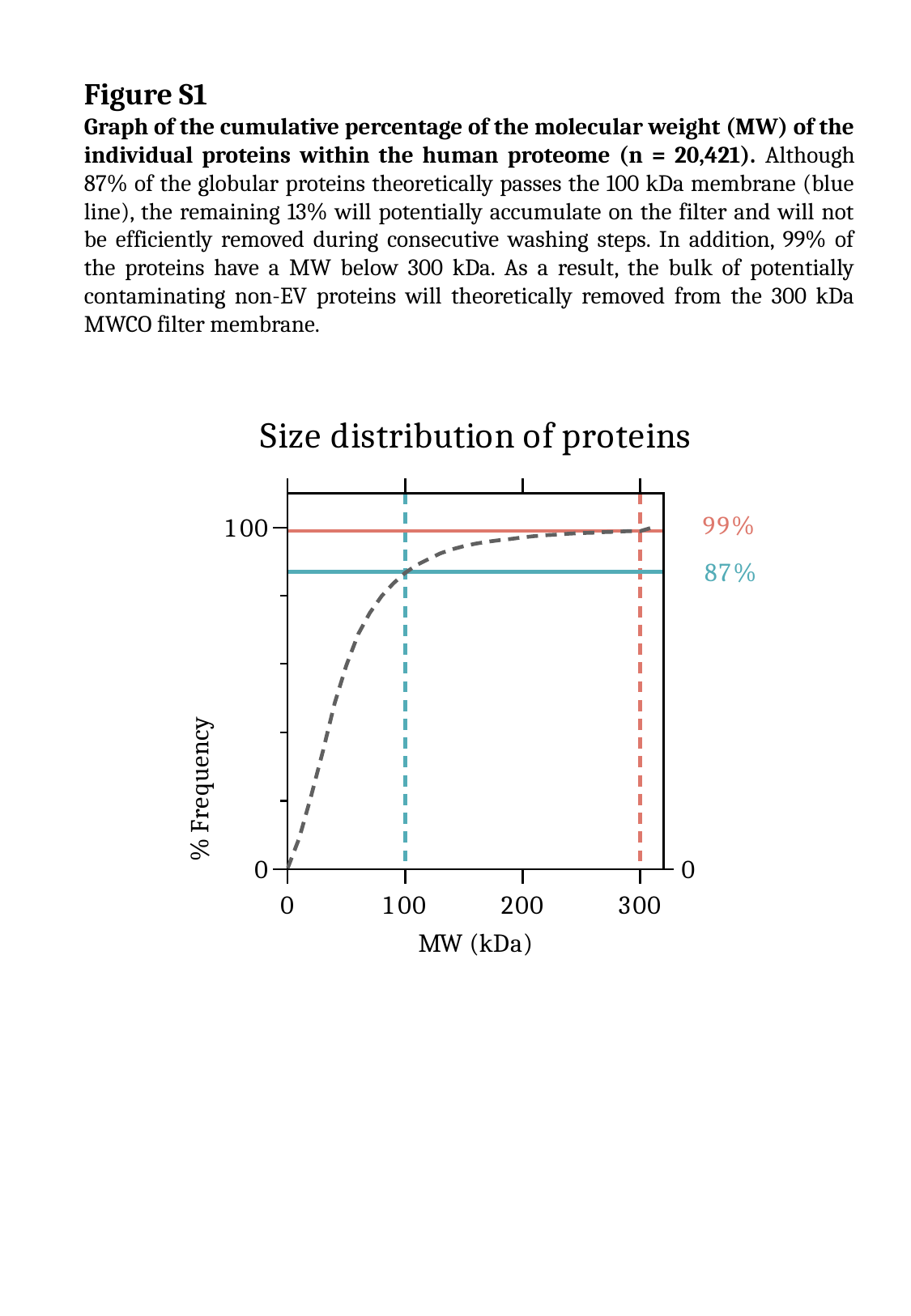

Figure S1
Graph of the cumulative percentage of the molecular weight (MW) of the individual proteins within the human proteome (n = 20,421). Although 87% of the globular proteins theoretically passes the 100 kDa membrane (blue line), the remaining 13% will potentially accumulate on the filter and will not be efficiently removed during consecutive washing steps. In addition, 99% of the proteins have a MW below 300 kDa. As a result, the bulk of potentially contaminating non-EV proteins will theoretically removed from the 300 kDa MWCO filter membrane.

### Slide 2
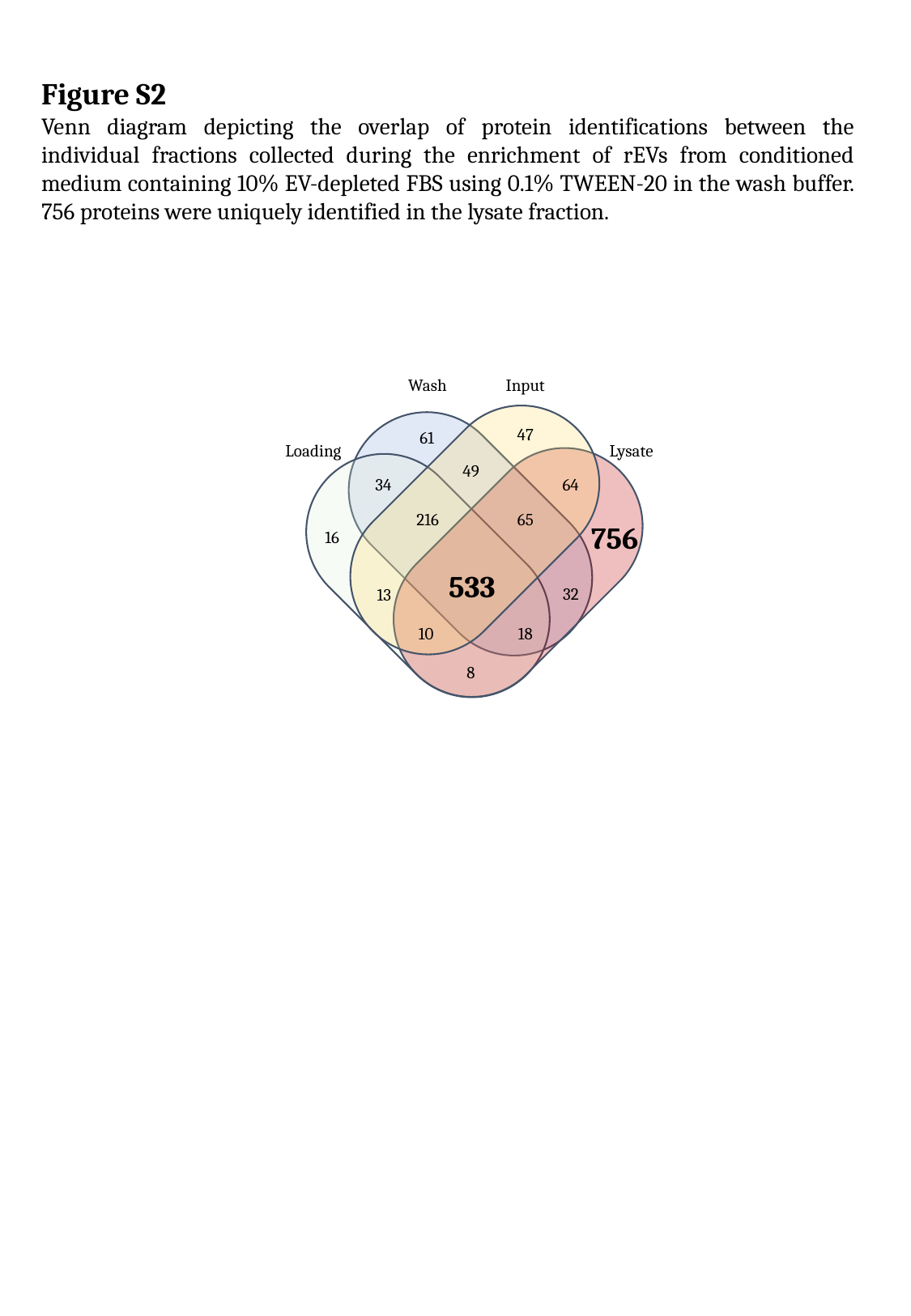

Figure S2
Venn diagram depicting the overlap of protein identifications between the individual fractions collected during the enrichment of rEVs from conditioned medium containing 10% EV-depleted FBS using 0.1% TWEEN-20 in the wash buffer. 756 proteins were uniquely identified in the lysate fraction.
Wash
Input
47
61
Loading
Lysate
49
34
64
216
65
756
16
533
32
13
18
10
8

### Slide 3
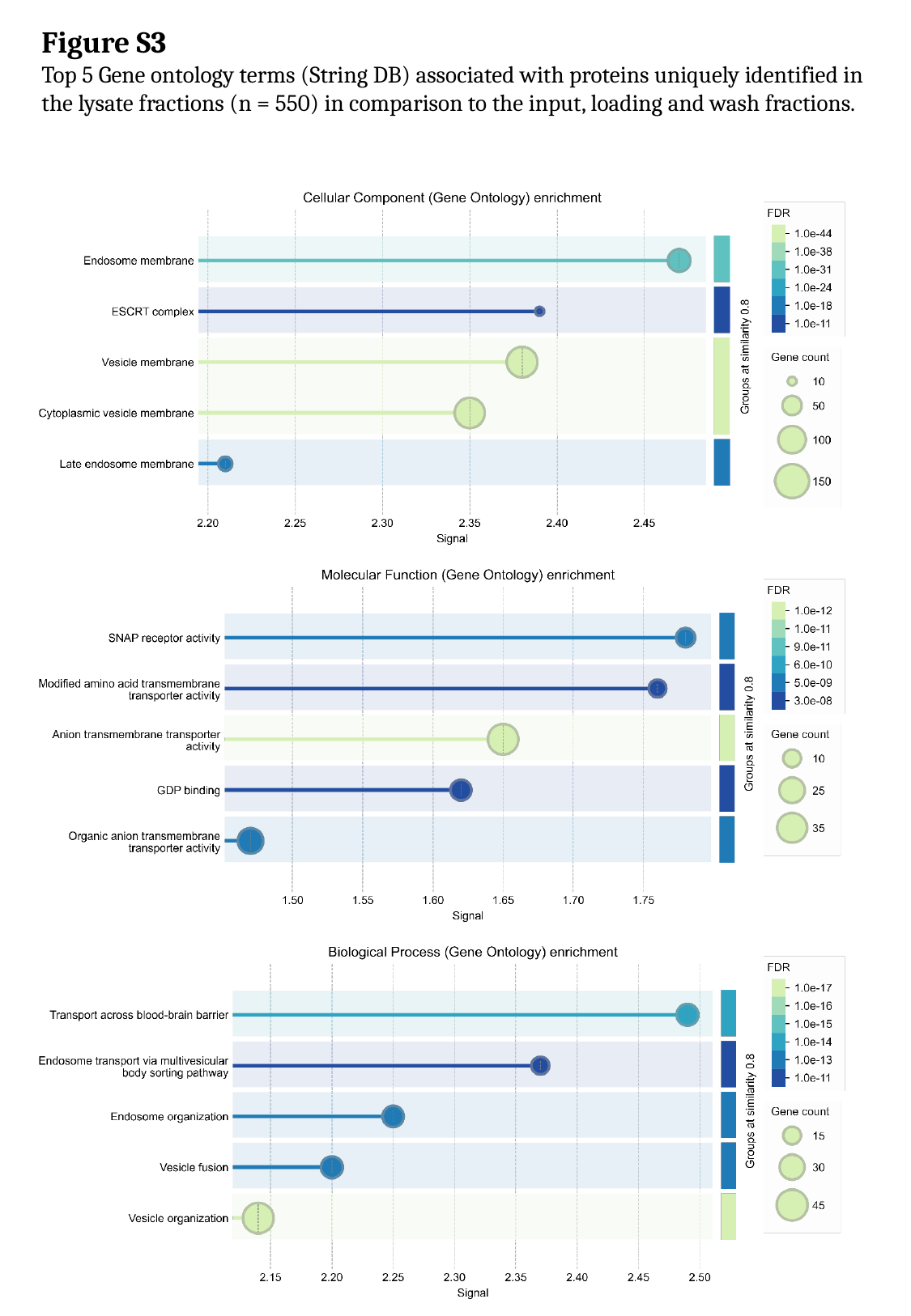

Figure S3
Top 5 Gene ontology terms (String DB) associated with proteins uniquely identified in the lysate fractions (n = 550) in comparison to the input, loading and wash fractions.

### Slide 4
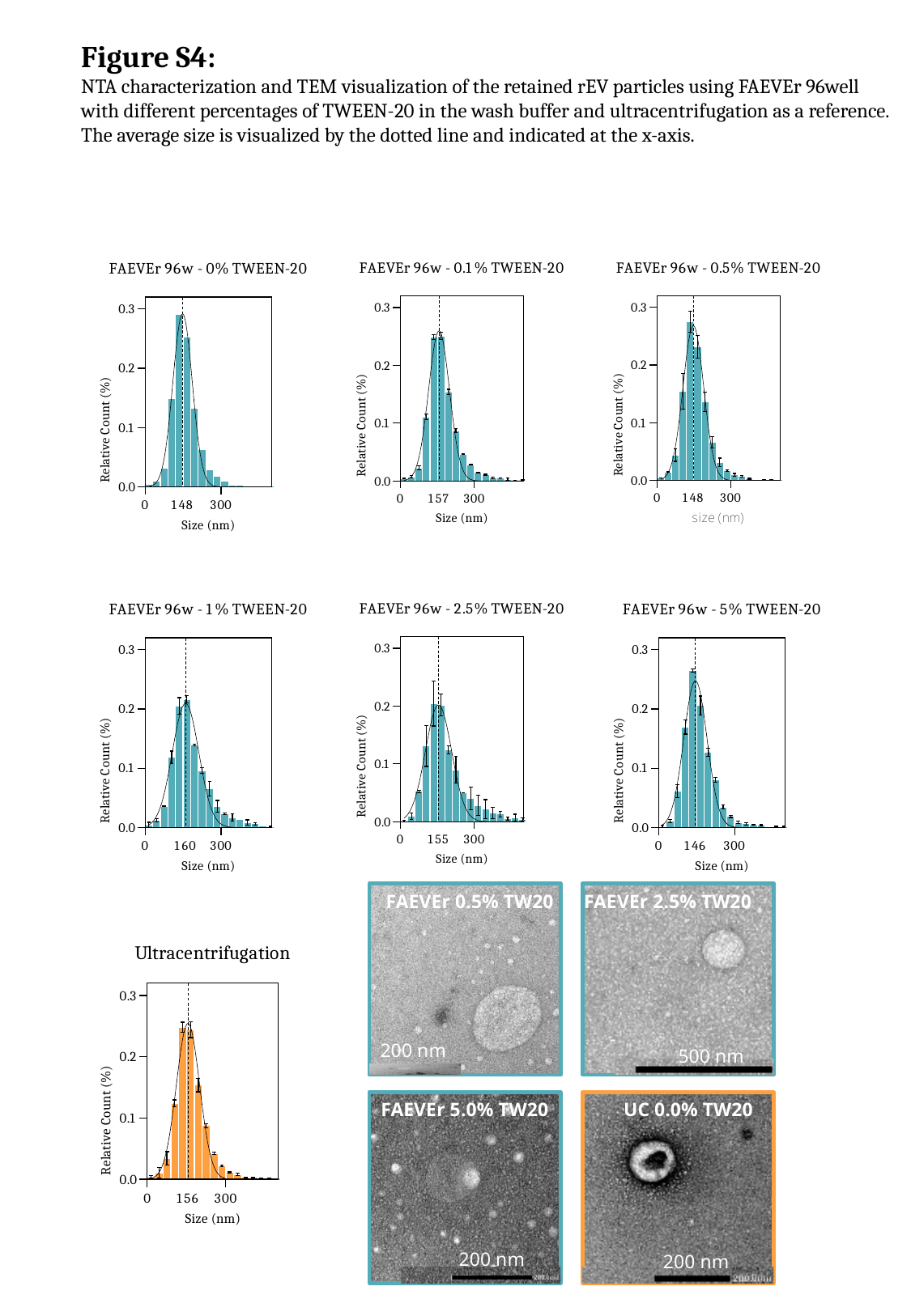

Figure S4:
NTA characterization and TEM visualization of the retained rEV particles using FAEVEr 96well with different percentages of TWEEN-20 in the wash buffer and ultracentrifugation as a reference. The average size is visualized by the dotted line and indicated at the x-axis.
200 nm
FAEVEr 0.5% TW20
FAEVEr 2.5% TW20
500 nm
200 nm
FAEVEr 5.0% TW20
200 nm
UC 0.0% TW20

### Slide 5
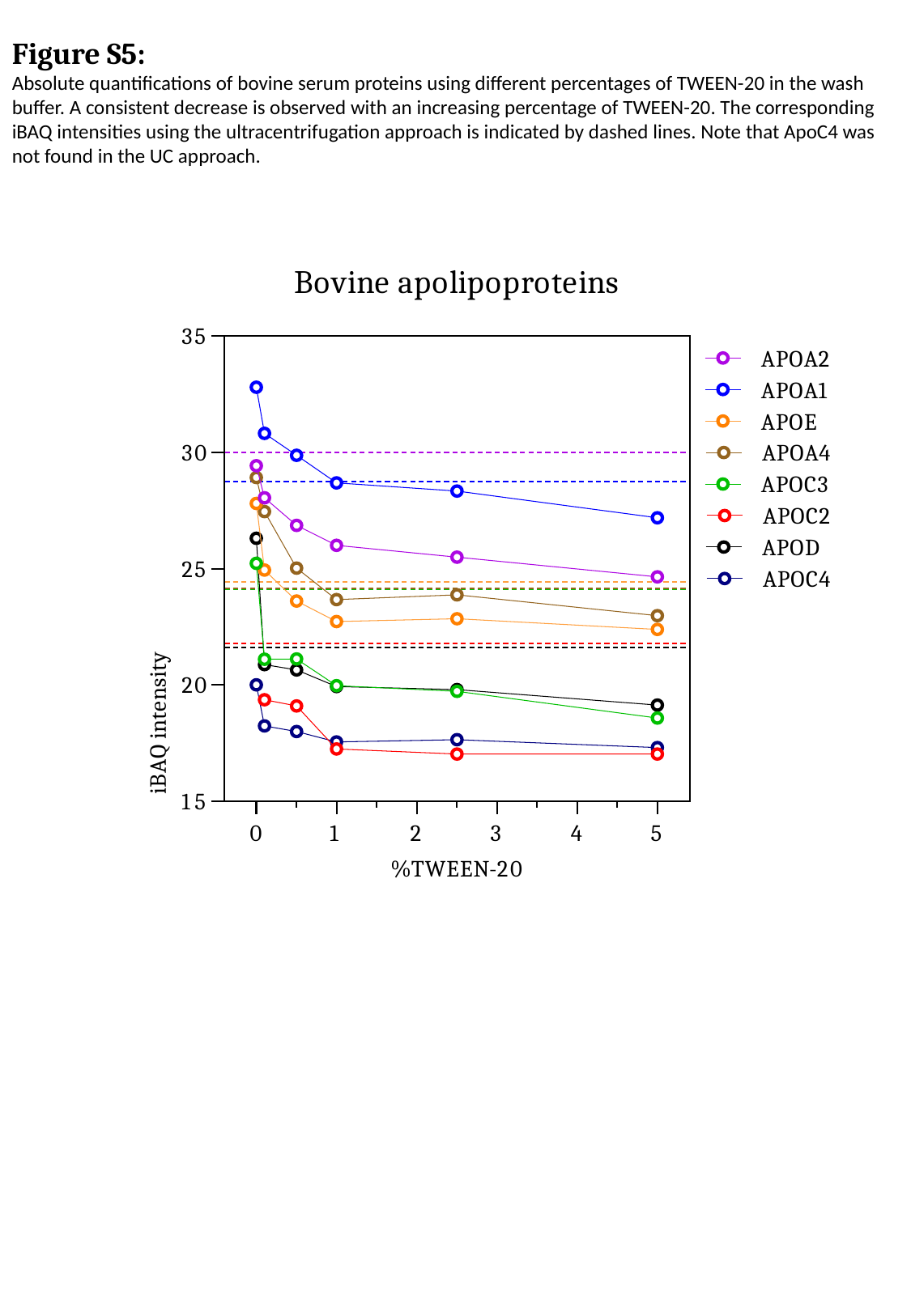

Figure S5:
Absolute quantifications of bovine serum proteins using different percentages of TWEEN-20 in the wash buffer. A consistent decrease is observed with an increasing percentage of TWEEN-20. The corresponding iBAQ intensities using the ultracentrifugation approach is indicated by dashed lines. Note that ApoC4 was not found in the UC approach.

### Slide 6
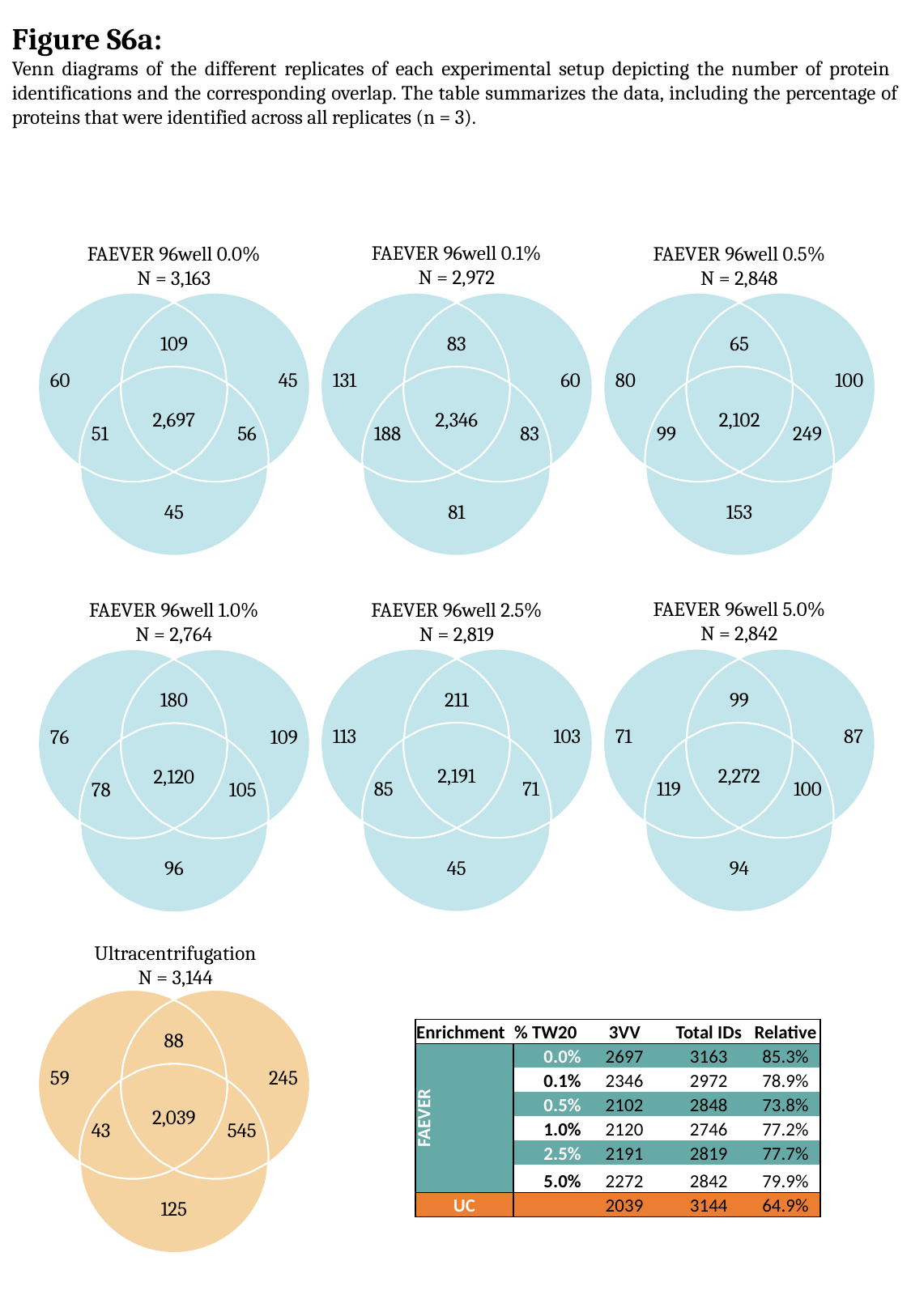

Figure S6a:
Venn diagrams of the different replicates of each experimental setup depicting the number of protein identifications and the corresponding overlap. The table summarizes the data, including the percentage of proteins that were identified across all replicates (n = 3).
FAEVER 96well 0.1%
N = 2,972
FAEVER 96well 0.5%
N = 2,848
FAEVER 96well 0.0%
N = 3,163
60
45
109
2,697
51
56
45
131
60
83
2,346
188
83
81
80
100
65
2,102
99
249
153
FAEVER 96well 5.0%
N = 2,842
FAEVER 96well 2.5%
N = 2,819
FAEVER 96well 1.0%
N = 2,764
113
103
211
2,191
85
71
45
71
87
99
2,272
119
100
94
76
109
180
2,120
78
105
96
Ultracentrifugation
N = 3,144
59
245
88
2,039
43
545
125
| Enrichment | % TW20 | 3VV | Total IDs | Relative |
| --- | --- | --- | --- | --- |
| FAEVER | 0.0% | 2697 | 3163 | 85.3% |
| | 0.1% | 2346 | 2972 | 78.9% |
| | 0.5% | 2102 | 2848 | 73.8% |
| | 1.0% | 2120 | 2746 | 77.2% |
| | 2.5% | 2191 | 2819 | 77.7% |
| | 5.0% | 2272 | 2842 | 79.9% |
| UC | | 2039 | 3144 | 64.9% |

### Slide 7
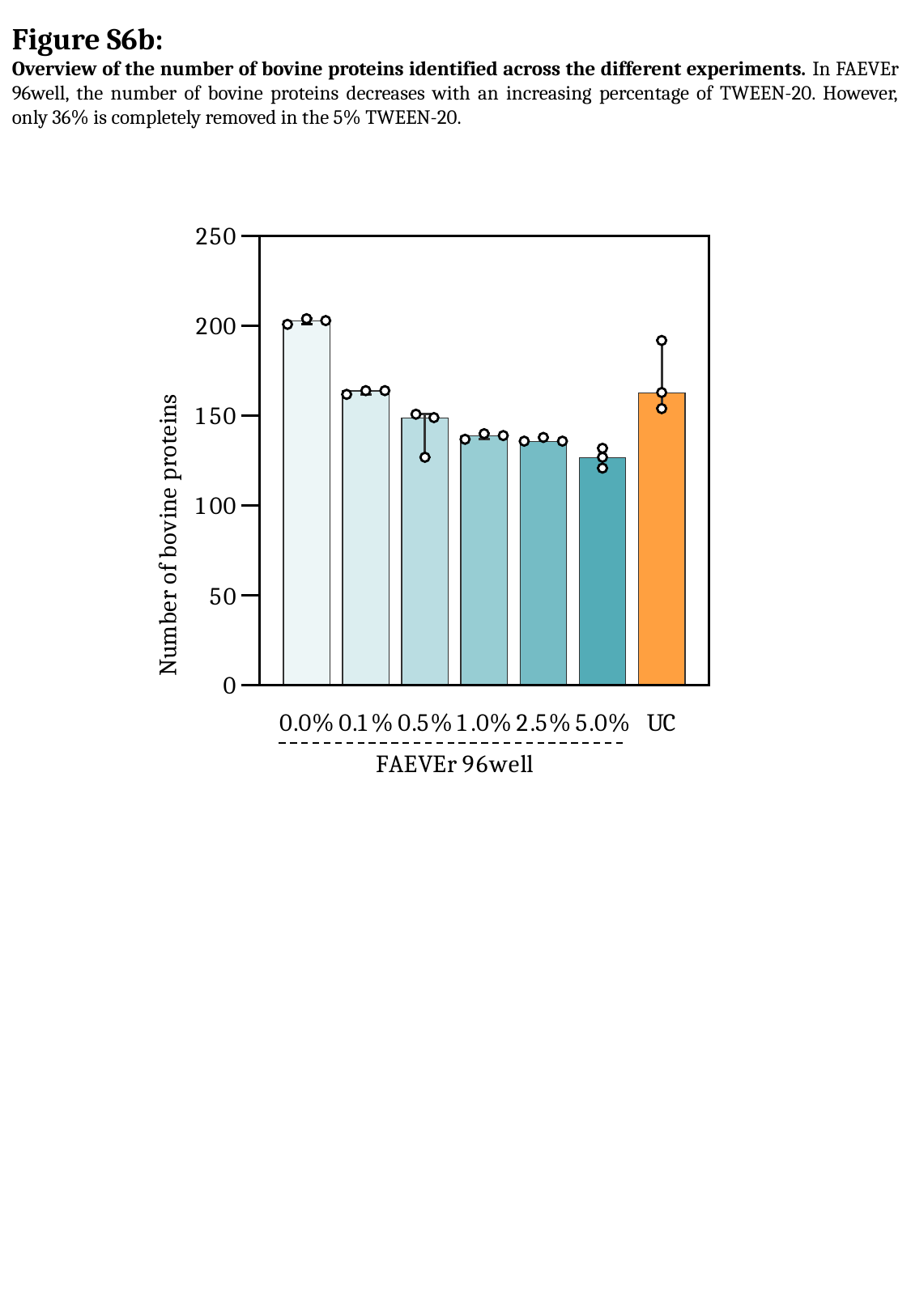

Figure S6b:
Overview of the number of bovine proteins identified across the different experiments. In FAEVEr 96well, the number of bovine proteins decreases with an increasing percentage of TWEEN-20. However, only 36% is completely removed in the 5% TWEEN-20.

### Slide 8
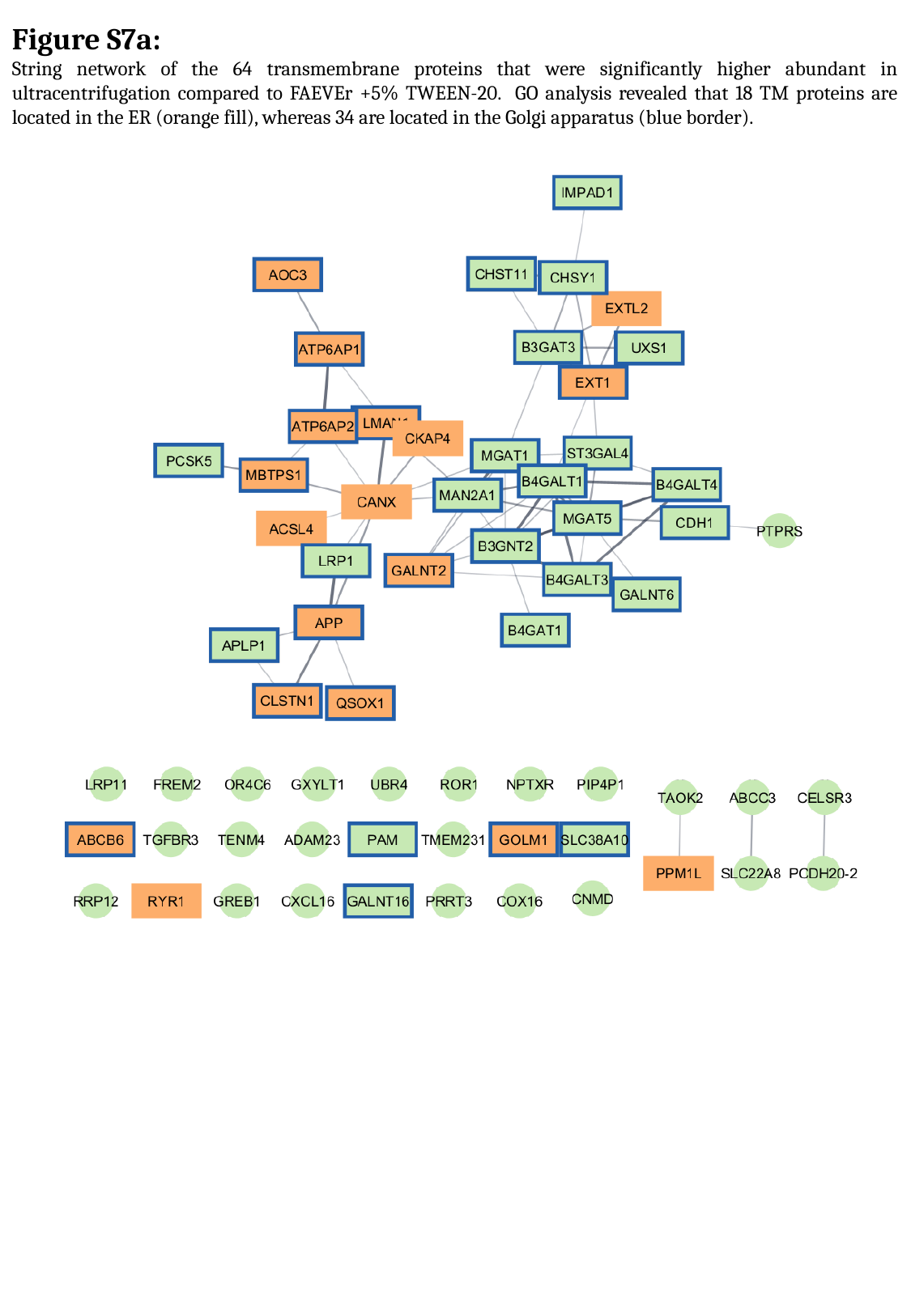

Figure S7a:
String network of the 64 transmembrane proteins that were significantly higher abundant in ultracentrifugation compared to FAEVEr +5% TWEEN-20. GO analysis revealed that 18 TM proteins are located in the ER (orange fill), whereas 34 are located in the Golgi apparatus (blue border).

### Slide 9
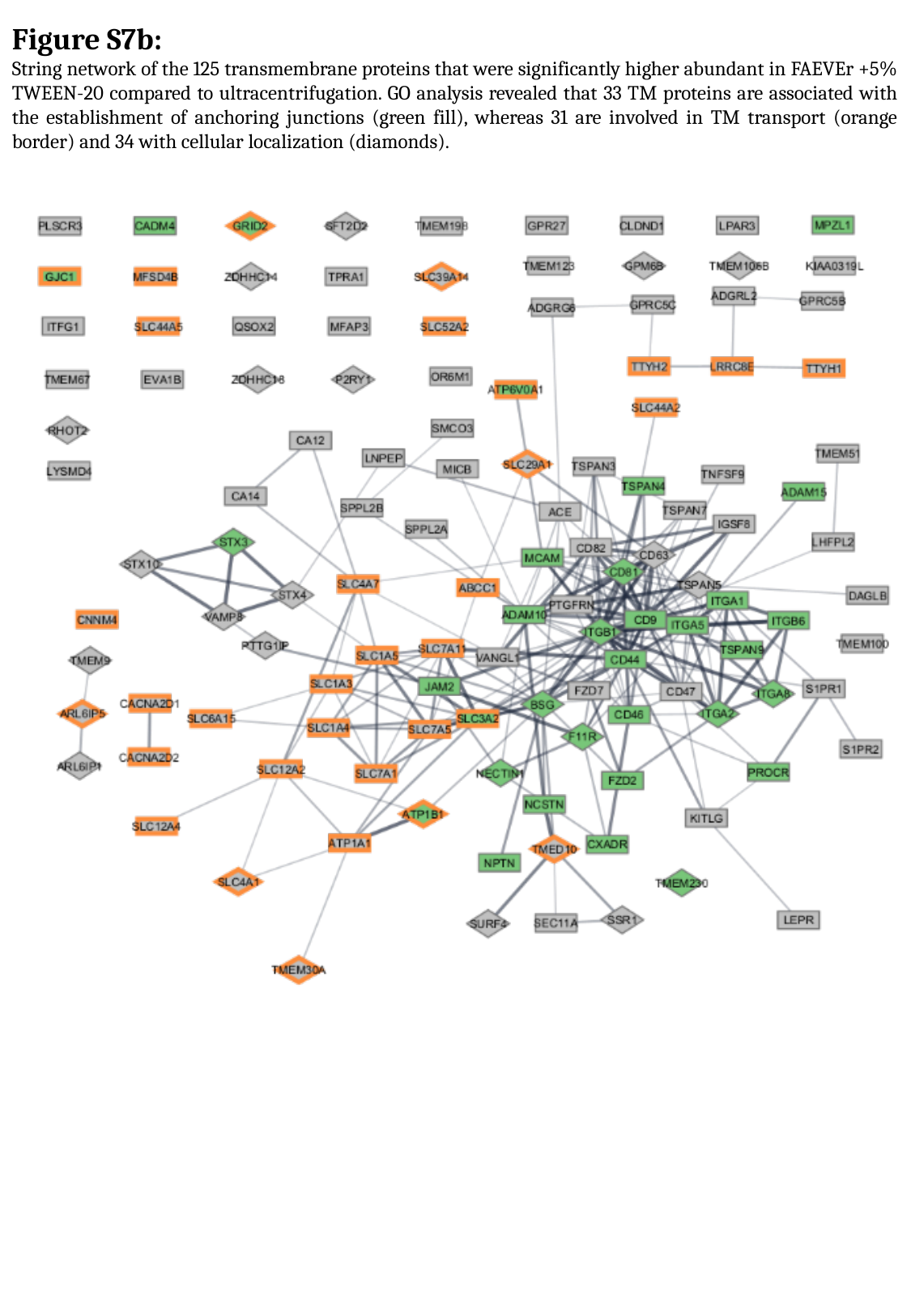

Figure S7b:
String network of the 125 transmembrane proteins that were significantly higher abundant in FAEVEr +5% TWEEN-20 compared to ultracentrifugation. GO analysis revealed that 33 TM proteins are associated with the establishment of anchoring junctions (green fill), whereas 31 are involved in TM transport (orange border) and 34 with cellular localization (diamonds).
